# Orange Maker: a CRISPR/Cas9-mediated genome editing and screening project to generate orange-eyed DarkJedi GAL4 lines by undergraduate students

**DOI:** 10.1101/2024.04.16.589788

**Authors:** Hee Su Park, Anna C. Gross, Seungjae Oh, Nam Chul Kim

## Abstract

One of the greatest strengths of *Drosophila* genetics is its easily observable and selectable phenotypic markers. The mini-white marker has been widely used as a transgenic marker for Drosophila transgenesis. Flies carrying a mini-white construct can exhibit various eye colors ranging from pale orange to intense red, depending on the insertion site and gene dosage. Because the two copies of the mini-white marker show a stronger orange color, this is often used for selecting progenies carrying two transgenes together in a single chromosome after chromosomal recombination. However, some GAL4 lines available in the fly community originally have very strong red eyes. Without employing another marker, such as GFP, generating a recombinant chromosome with the strong red-eyed GAL4 and a desired UAS-transgene construct may be difficult. Therefore, we decided to change the red eyes of GAL4 lines to orange color. To change the eye color of the fly, we tested the CRISPR/Cas9 method with a guide RNA targeting the white gene with OK371-GAL4 and elav-GAL4. After a simple screening, we have successfully obtained multiple lines of orange-eyed OK371-GAL4 and elav-GAL4 that still maintain their original expression patterns. All of these simple experiments were performed by undergraduate students, allowing them to learn about a variety of different genetic experiments and genome editing while contributing to the fly research community by creating fruit fly lines that will be used in real-world research.

## Introduction

*Drosophila melanogaster*, the common fruit fly (or vinegar fly), has been a pivotal model organism in the field of genetic research for over a century, offering profound insights into the mechanisms of genetic inheritance, development, neurobiology, and human diseases^1^. Its compact genome, short life cycle, and the ease with which it can be manipulated genetically make it an invaluable tool for understanding complex biological processes. One key feature that has facilitated its wide use in genetic experiments is the presence of easily observable phenotypic markers, such as eye color. These markers have been instrumental in the development of sophisticated genetic tools like the GAL4/UAS system^2^, which allows for targeted gene expression in specific tissues or at particular developmental stages. Ever since the development of the GAL4/UAS system, it has become an indispensable tool for expressing exogenous genes in *Drosophila melanogaster* to study their function and identify associated genetic interactors. The GAL4/UAS system involves a yeast-derived transcription factor (GAL4) that binds to upstream activating sequences (UAS) to control the expression of downstream genes^2^. Notably, the GAL4/UAS system in *Drosophila* is a bipartite system composed of a tissue-specifically expressed GAL4 and a transgene under the control of UAS^2^. Therefore, GAL4 driver lines and UAS-transgene lines can be separately generated and maintained. Crossing two parental lines carrying a GAL4 and a UAS-transgene allows transgene expression in their progenies^2^. This bipartite system essentially increases the efficiency of developing toolkits such as tissue- or cell-specifically expressed GAL4 driver lines and whole-genome level transgenic tools such as RNAi ^1^.

To generate transgenic lines of GAL4 or UAS-transgene, the mini-white gene has been widely used as a marker aiding in identifying successful gene insertions through the observable phenotype of eye color variation. The mini-white gene is a derivative of the white gene^3^, which is naturally responsible for the eye color in fruit flies. The wild-type white gene leads to red eyes, while mutations in the white gene can result in white eyes due to a lack of pigmentation. The white gene encodes ATP-binding cassette G-subfamily (ABCG) half-transporter transporting precursors required for red pigments^4^. When host flies with white eyes are successfully engineered with a transgenic plasmid construct carrying the mini-white gene, the transgenic animals can have various eye colors ranging from very light-yellowish orange color to strong red. This eye color variation with the mini-white gene depends on transcriptional gene expression activity of the location where the transgenic construct is inserted and how many mini-white genes they have^5^. When the mini-white is located in a transcriptionally active area, or flies have multiple copies of mini-white, their eye color will be more intense and closer to red.

Despite its widespread application, the mini-white marker system faces challenges when utilized in conjunction with certain GAL4 lines that exhibit intense red eyes. This presents a substantial hurdle in the generation and selection of recombinant chromosomes, especially when aiming to pair a strong red-eyed GAL4 line with a specific UAS-transgene construct. The overlap in eye color phenotypes renders the task of distinguishing between the parental and recombinant flies imprecise without the aid of additional markers. Addressing this limitation, our study leverages the specificity and efficiency of the CRISPR/Cas9 gene-editing technology to modify the eye color expression in selected GAL4 lines by mutating the mini-white gene to make it less active. By employing a previously published guide RNA targeted towards the white gene in conjunction with germline-specifically expressed Cas9^6,7^, we aim to shift the eye color from red to orange, thus enabling a more accurate selection of recombinant progenies without any additional markers.

This report chronicles our experimentation with two specific GAL4 lines—OK371-GAL4 and elav-GAL4—both of which inherently exhibit strong red eye pigmentation. Through the utilization of CRISPR/Cas9-mediated gene editing, we have succeeded in producing multiple orange-eyed lines for both GAL4 strains. These modified lines offer a distinct phenotype that eases the selection process in genetic crosses. We conducted in-depth analyses to ensure that the alterations have not compromised the GAL4 activity, thereby preserving the integrity of these vital genetic tools for future research endeavors. More importantly, undergraduate students performed these experiments without specialized equipment except the confocal microscope used for examining precise expression patterns. Through this simple experimental system, any undergraduate or high school student can gain knowledge and hands-on experience with a variety of different genetic experiments and genome editing while contributing to the fly research community by creating fruit fly lines that will be used in real-world research.

## Main Text Methods

### Drosophila Lines

Guide RNAs: y2 cho2 v1; attP40{U6.2-w-ex3-2} (NIG-FLY stock #: GRN-002) Cas9: y[1] M{w[+mC]=nos-Cas9.P}ZH-2A w[*] (Bloomington stock #: 54591) Elav^C155^-GAL4: P{w[+mW.hs]=GawB}elav[C155] (Bloomington stock #: 458) OK371: w[1118]; P{w[+mW.hs]=GawB}VGlut[OK371] (Bloomington stock #: 26160) JFRC81-GFP was obtained from the Rubin laboratory(ref). Balanced Cas9: y[1] M{w[+mC]=nos-Cas9.P}ZH-2A w[*];Bl/CyO (This study) Founder line (Orange Maker): y[1] M{w[+mC]=nos-Cas9.P}ZH-2A w[*];attP40{U6.2-w-ex3-2}/CyO (This study, white eyed)

### Fly Maintenance and crosses

Flies were crossed, grown on standard Bloomington Drosophila Stock Center cornmeal food, and maintained at 25 °C with a standard 12h:12h light/dark cycle.

#### Balanced Cas9 generation

To generate the founder line expressing Cas9 and guide RNAs in germline cells, we first generated the balanced Cas9 line by crossing virgin female Cas9 (#54591) with male Bl/CyO flies. Male flies carrying Cas9 and CyO, or Cas9 and Bl, were mated with virgin Cas9 females again. Then, virgin females carrying Cas9 and CyO were collected and mated with male Cas9 and Bl. Finally, virgin females carrying Cas9, Bl, and CyO were collected and mated with male flies carrying Cas9, Bl, and CyO (Fig. S1A).

#### Founder line (Orange Maker) generation

Virgin female flies of the balanced Cas9 (y[1] M{w[+mC]=nos-Cas9.P}ZH-2A w[*]; Bl/CyO) were crossed with males of GRN-002 (y2 cho2 v1; attP40{U6.2-w-ex3-2}). Male flies carrying Cas9 and GRN-002 balanced with CyO were collected and crossed with virgin females of the balanced Cas9 again. Virgin females carrying Cas9 and GRN-002 balanced with CyO were collected and mated with male files carrying Cas9 and GRN-002 balanced with CyO (Fig. S1B).

#### Orange Making Crosses (DarkJedi line generation)

Virgin female flies of the founder line (y[1] M{w[+mC]=nos-Cas9.P}ZH-2A w[*];attP40{U6.2-w-ex3-2}/CyO) were collected and mated with males of OK371-GAL4 (Fig. S1C) or elav^C155^-GAL4 (Fig. S1D). In the case of OK371, male flies with mosaic eyes carrying Cas9, GRN-002, and OK371* (modified) were collected, and individual males were crossed with virgin female Bl/CyO flies. Orange-eyed males carrying modified versions of OK371 were individually mated with virgin female Bl/CyO flies. Virgin females and males with orange color eyes and CyO were crossed to establish a OK371^DJ^ line. In the case of elav^C155^-GAL4, after the first cross, female virgin flies with mosaic eyes carrying Cas9, GRN-002, and elav^C155^-GAL4* (modified) were collected, and individual females were crossed with male w^1118^. In order to simplify the cross and to keep up with the OK371 experiment, GRN-002 was left in a floating state. Male flies with orange eyes (elav^C155DJ^) were collected and mated with female virgin w^1118^ flies. Then, orange-eyed males and females were collected and mated. Finally, homozygous virgin female flies were collected by their stronger orange eyes and crossed with orange-eyed males (These lines may still have GRN-002 floating, Fig. S1D).

### Imaging

*Drosophila* eye pictures were taken using a Leica Z16 APO with the DMC2900 camera and a ring light. Larval images without dissection were taken by a stereoscope (Amscope, SF-2TR-WF) equipped with a digital camera (Amscope, MU1000), a filter (Edmunds optics, 66-085), and a blue LED light (ebay, PAR30). Fluorescent larval brain images were taken using a Nikon X-light2 spinning disc confocal microscope. Larval brains were dissected in PBS and fixed with 4% paraformaldehyde for 10 minutes at room temperature. The fixed larval brains were mounted on the slide glass with Fluoromount-G (Southern biotech). The Z-stack images were obtained with a Nikon Plan APO 40x SIL silicone immersion objective (NA 1.25). Maximum Z-projection images were generated by ImageJ (NIH).

## Results

To make red eyes of OK371 and elav^C155^-GAL4 by introducing mutations using the CRISPR/cas9 system, we first generated a founder line expressing sgRNA targeting the white gene in the germline cells (Fig. 1, see methods and Fig. S1 for detailed mating schemes). In this combination founder line (briefly, nos-Cas9; GRN-002/CyO), as they express Cas9 germline-specifically with nos promoter and a guide RNA with U6 promoter, their eye color became white due to mutations in the mini-white marker in the Cas9 construct. The GRN-002 targets the third exon of the white gene with a high mutagenesis efficiency ^6^. Although high mutation frequencies were observed in the male germline ^6^, we decided to use the female germline. Maternally deposited Cas9 will generate somatic mosaic mutants, which will excite students. Thus, female virgins of the founder line were crossed with male OK731 or elav^C155^-GAL4 (Fig. 1 and methods). Progenies of these crosses were predominantly white-eyed. However, flies with mosaic eyes were produced occasionally (Fig. S2A). Male mosaic-eyed animals were collected and used for crosses to produce orange-eyed flies. Subsequently, male flies having pale orange eye colors were collected and individually crossed to make stable lines (see methods and Fig. S1C and D for detailed mating schemes). Finally, we successfully generated multiple orange-eyed GAL lines from originally red-eyed OK371 and elav^C155^-GAL4 lines. These lines have various eye colors for OK371(Fig. 2) and elav^C155^-GAL4 (Fig. 3). We have designated these variant GAL4 lines generated by the Orange Maker as the DarkJedi (DJ) lines, inspired by the orange lightsabers of the Dark Jedi, which contrast with the red lightsabers of the Sith in the Star Wars series. In both OK371^DJ^ and elav^C155DJ^ lines, heterozygous males exhibit stronger color than the color of heterozygous females (Fig. 2A and 3A). In the case of homozygous flies, some lines show a red color as strong as the original lines (Fig. 2B and 3B). Overall, we got more diverse color variants in OK371 as we established more lines (Fig 2 and 3).

**Figure 1.**
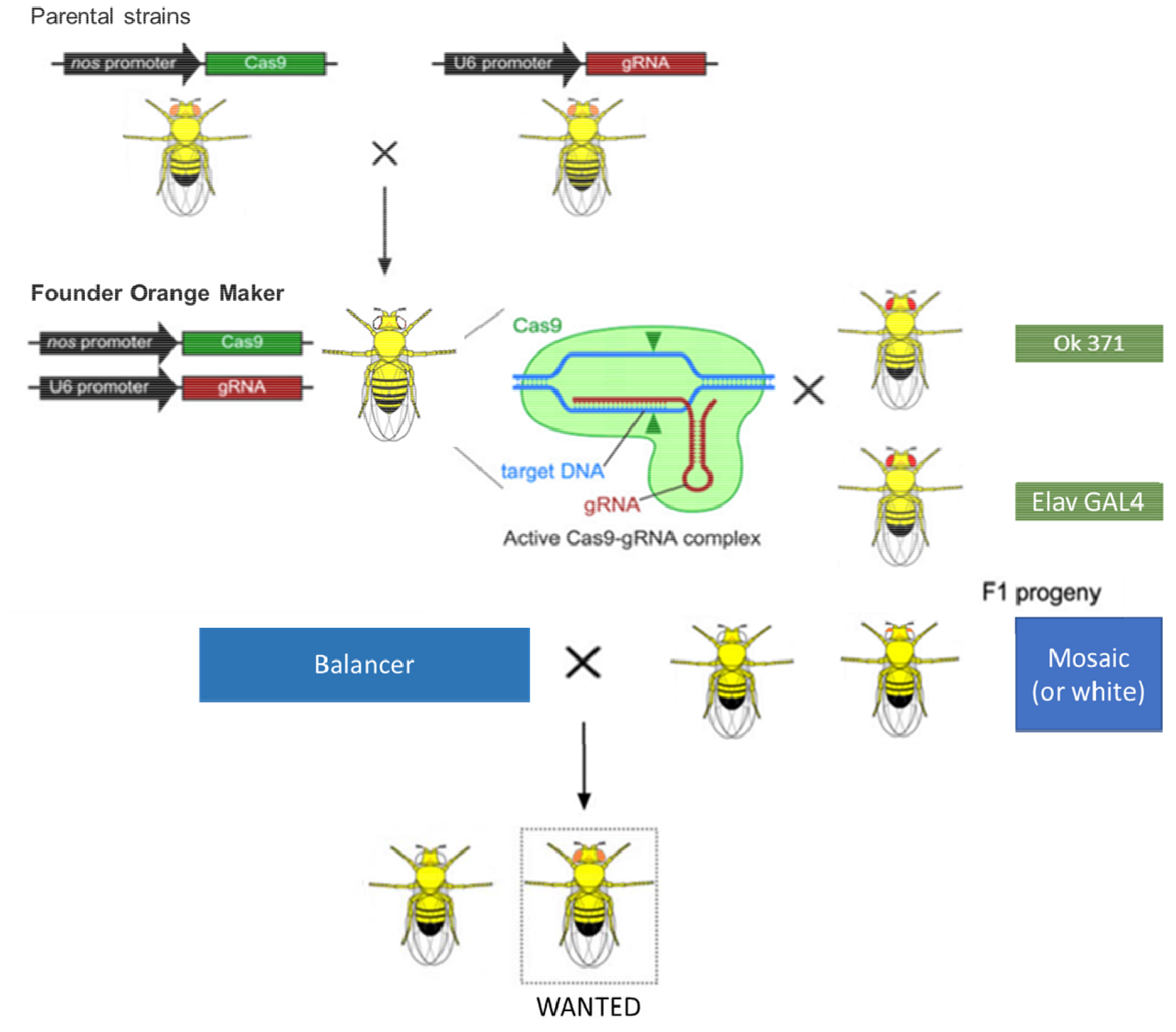
Schematic diagram of orange making. Using two previously published two parental lines, we generated a founder Orange Maker carrying both germline-expressed Cas9 and synthetic guide-RNA targeting the white gene. Then, females of Orange Maker were used to generate flies having mosaic eyes. Although both mosaic and white-eyes flies can be used for orange making, we used only mosaic eyed flies (see methods and supplementary figure 1 for detailed mating schemes, modified from Kondo and Ueda, 2013).

**Figure 2.**
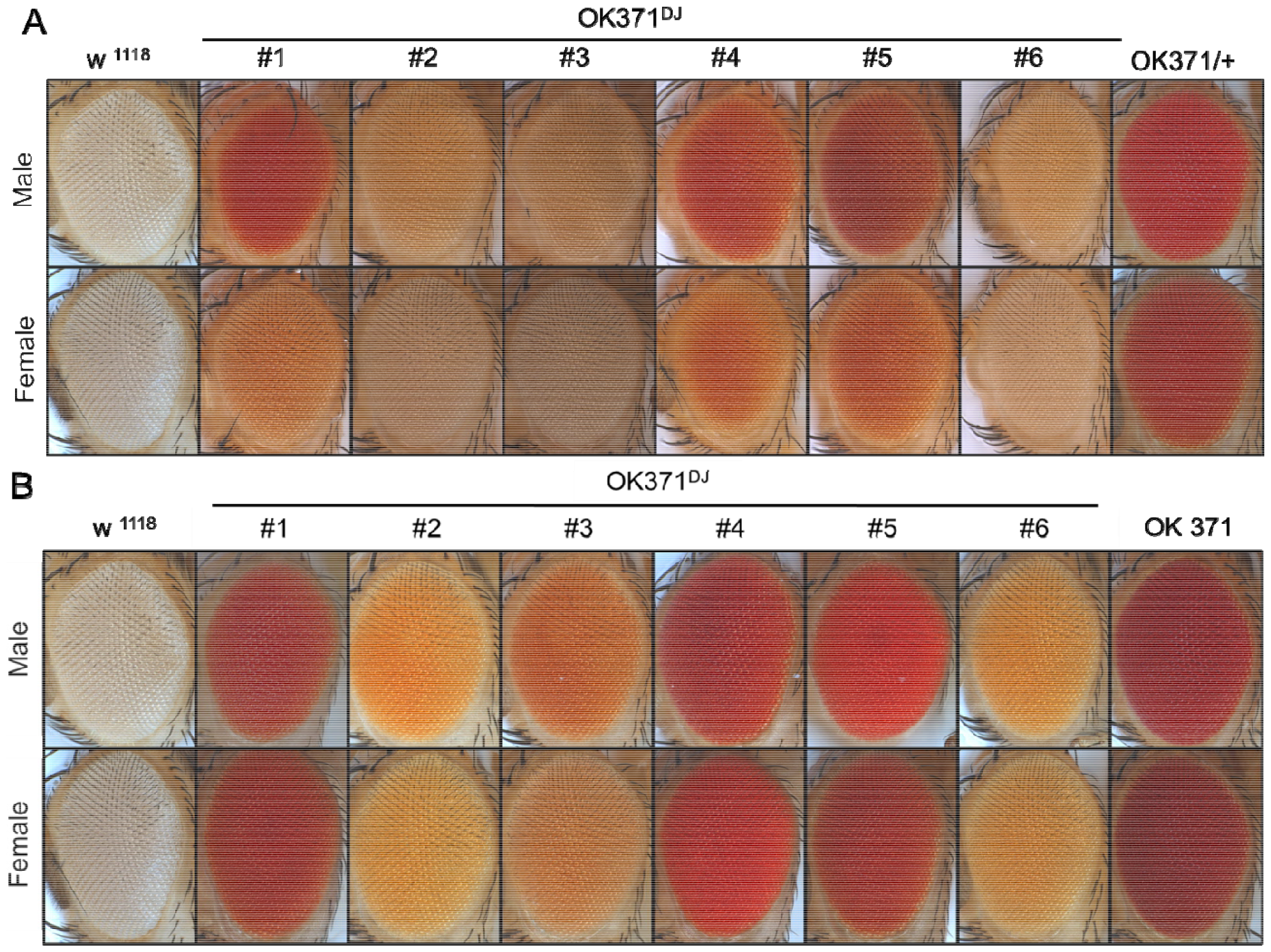
Red Or Orange. Various eye colors generated by targeting the mini-white gene in OK371. (A) Eye colors of heterozygous OK371 DarkJedi (DJ) lines, (B) Eye colors of homozygous OK371 DarkJedi (DJ) lines. White-eyed w^1118^ and red-eyed original OK371 were used as controls.

**Figure 3.**
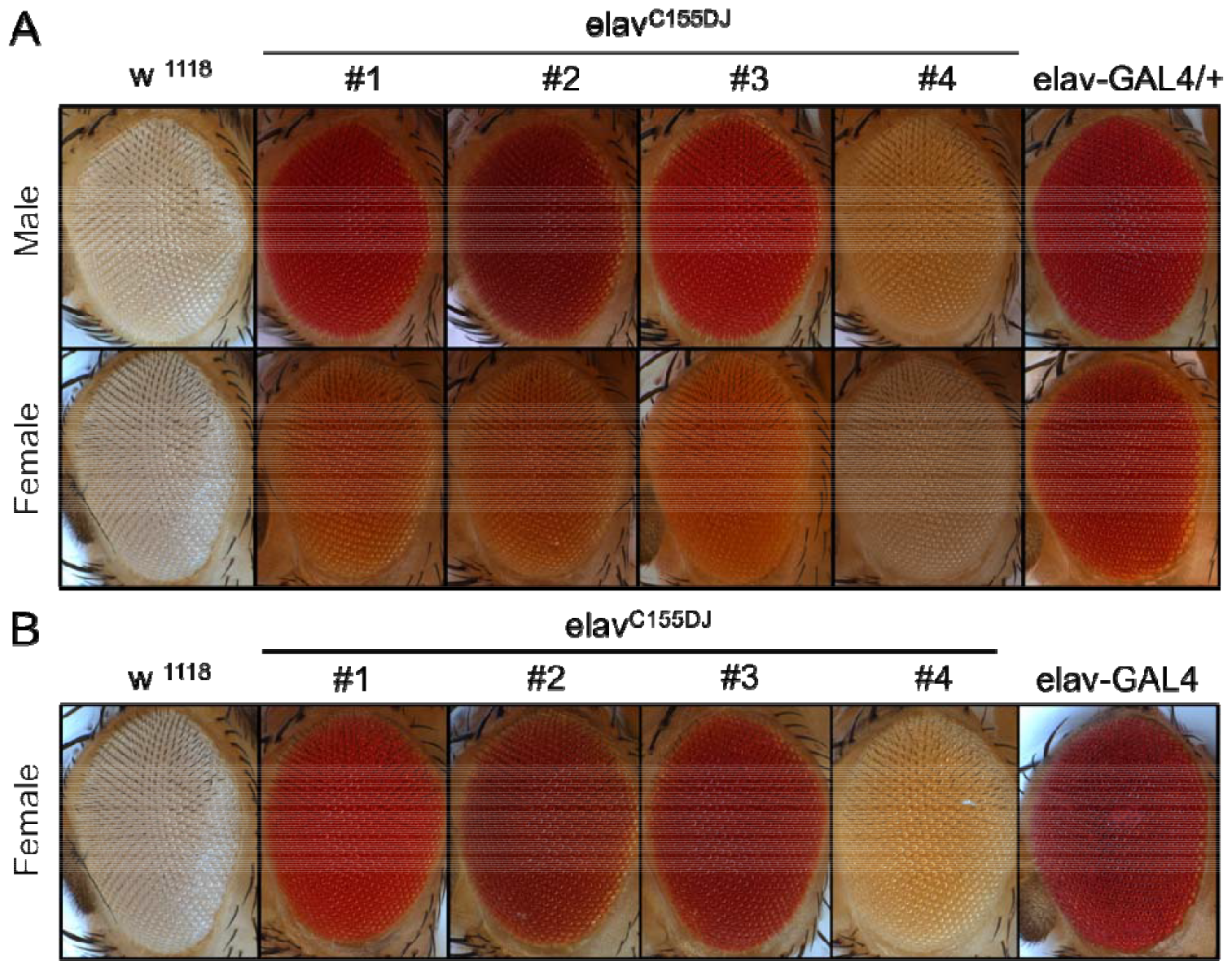
Red Or Orange. Various eye colors generated by targeting the mini-white gene in elav^C155^-GAL4. (A) Eye colors of hemizygous male and heterozygous female elav^C155^ DarkJedi (DJ) lines, (B) Eye colors of homozygous female elav^C155^ DarkJedi (DJ) lines. White-eyed w^1118^ and red-eyed original elav^C155^-GAL4 were used as controls.

To examine whether these modified versions of GAL4 lines have similar gene expression patterns and strength, we crossed newly established GAL4 lines with JFRC81-GFP ^8^. GFP expression with JFRC81-GFP, OK371, and elavC155-GAL4 lines was strong enough to be detected in salivary glands and ventral ganglions without dissection by our DIY fluorescence stereoscope setup (methods and Fig. S2B and C). Therefore, we further examined their expression with a confocal microscope. Most of these lines expressed comparable levels of GFP, and the expression patterns were generally similar to the original expression patterns (Fig. 4A and B). Although we did not sequence these flies, this result implies that targeting the mini-white maker gene might only introduce local ins/del mutations that did not affect nearby GAL4 expression.

**Figure 4.**
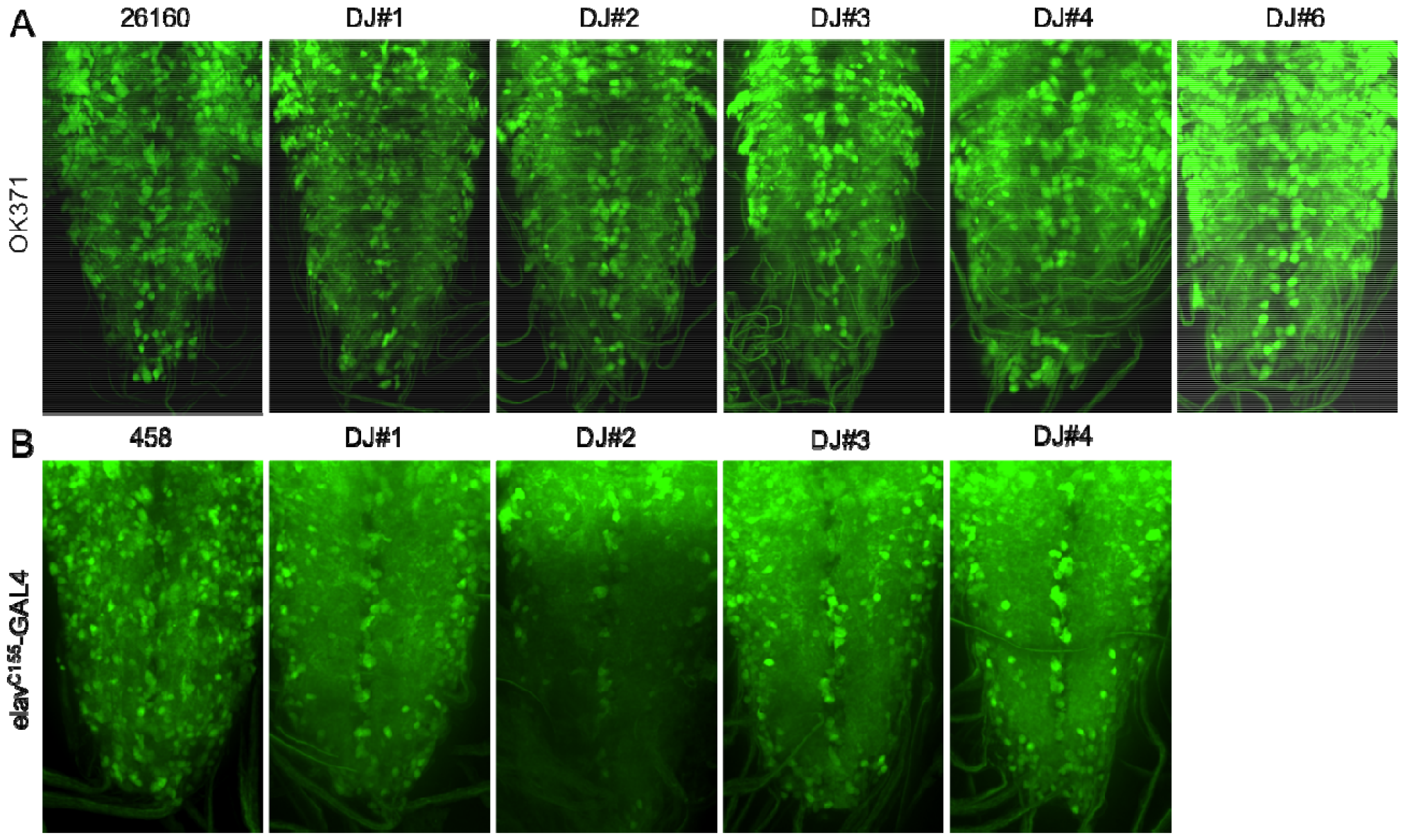
GFP expression patterns derived by OK371^DJ^ and elav^C155DJ^ lines in the *Drosophila* larval ventral ganglion. (A) GFP expression with OK371^DJ^ lines, (B) GFP expression with elav^C155DJ^ lines. Original stocks for OK371 (BL 26160) and elav^C155^-GAL4 (BL 458) were used as controls.

Through this orange maker experiment, undergraduate students have learned about *Drosophila* transgenesis, the GAL4/UAS system, mendelian genetics, somatic mosaicism, and the CRISPR/Cas9 system. In addition, we have successfully generated new GAL4 lines exhibiting various eye colors that can be used in the fly research community. OK371^DJ^ lines can be readily used for functional experiments as we have isolated the second chromosome carrying variant OK371 while establishing the lines.

## Discussion

This study had two purposes. Educationally, this study was designed to provide an opportunity for undergraduate students or advanced high school students to learn about genetics and genome editing. Scientifically, this study aimed to generate orange-eyed versions of traditional GAL4 lines for an efficient recombinant chromosome selection with two transgenes. More importantly, through this study, undergraduate authors not only learned about this system but also generated *Drosophila* lines that can be used in the fly research community. As we have designated variant GAL4 lines generated by the Orange Maker as the DarkJedi lines and marked by superscript DJ and serial numbers on the original GAL4 names, other variant GAL4 lines generated by other students can also be named as the DarkJedi lines and be easily distinguishable from their original liens by eye colors and names.

However, to be used in real research, these lines should be further validated by genomic sequencing. Genomic sequencing may reveal the relationship between eye colors and various mutations on the mini-white gene. In general, the size of deletion or insertion is expected to be inversely related to the intensity of eye color. If necessary, mutant mini-white genes may be used as a new variant marker for *Drosophila* transgenesis. In addition, these lines may need to be backcrossed with w^1118^ several times to remove unexpected events that happened during the orange-making processes. Especially, elav^C155^-GAL4 lines may still have unnecessary guide-RNA, and these lines may also have unexpected genome rearrangement because the endogenous white gene and the mini-white on the X chromosome were targeted.

## Acknowledgments

We thank the Bloomington *Drosophila* Stock Center and NIG-Fly for the *Drosophila* stocks. We also thank the University of Minnesota Summer College of Pharmacy Experience (SCoPE) Program.

## Author contributions

N.C.K. conceived the project. A.C.G. and N.C.K. initiated and conducted all the genetic experiments. H.S.P. and S.O. conducted genetic experiments and performed all image analyses. N.C.K. and H.S.P. wrote the manuscript. N.C.K. supervised the project.

## Funding

Financial support was provided by the University of Minnesota Summer College of Pharmacy Experience (SCoPE) Program to Anna Gross.

## Declarations

### Ethics approval and consent to participate

Not applicable.

### Availability of data and material

The data are available from the corresponding author upon request.

### Consent for publication

Not applicable.

### Competing interests

None.

---

**Supplementary Figure 1.**
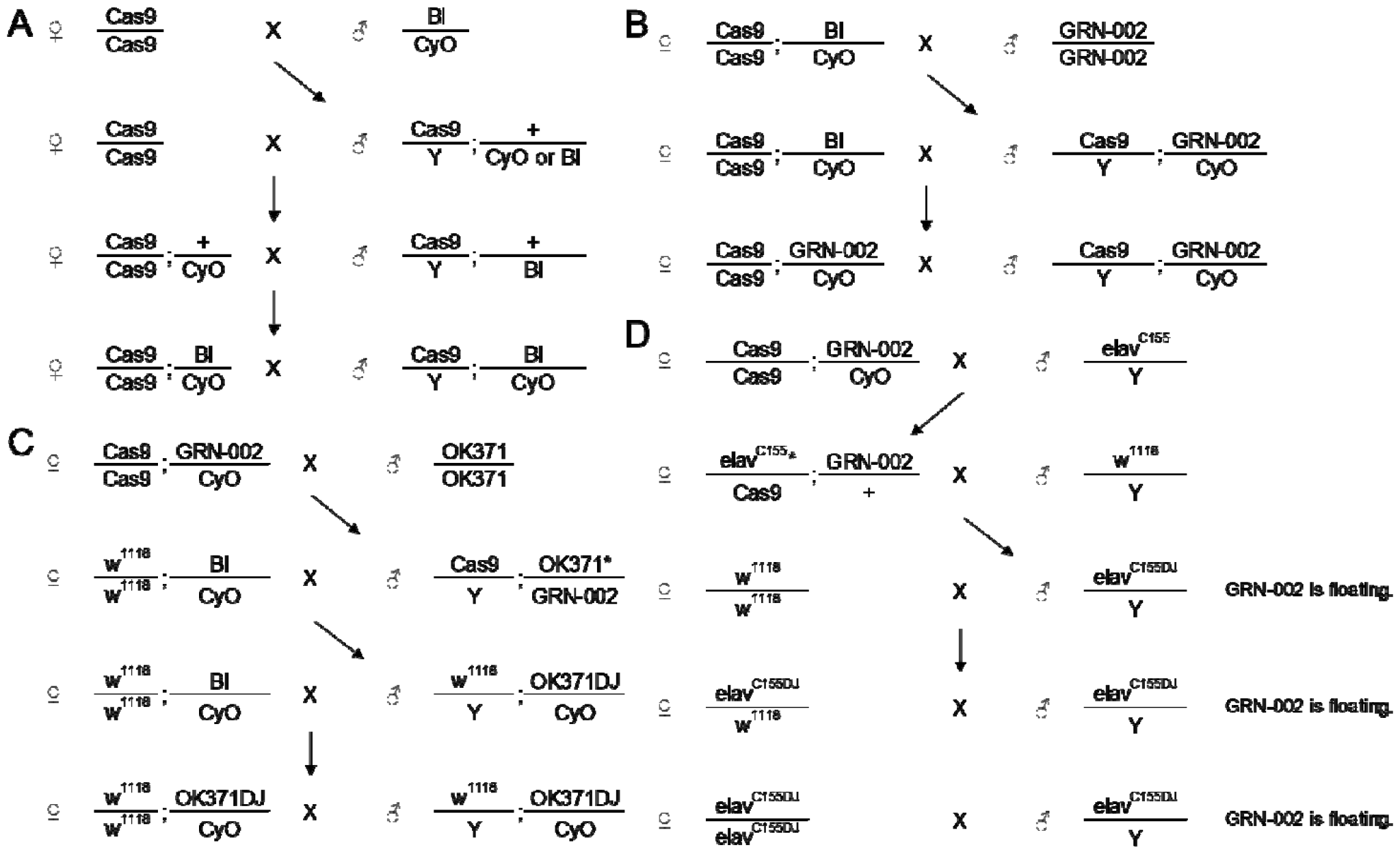
Mating schemes for the generation of the Orange Maker and orange making with the Orange Maker. (A) Balanced Cas9 generation, (B) Orange Maker generation, (C) Orange making (DarkJedi line generation) with OK371, (D) Orange making (DarkJedi line generation) with elav^C155^-GAL4, see the method section for a detailed explanation.

**Supplementary Figure 2.**
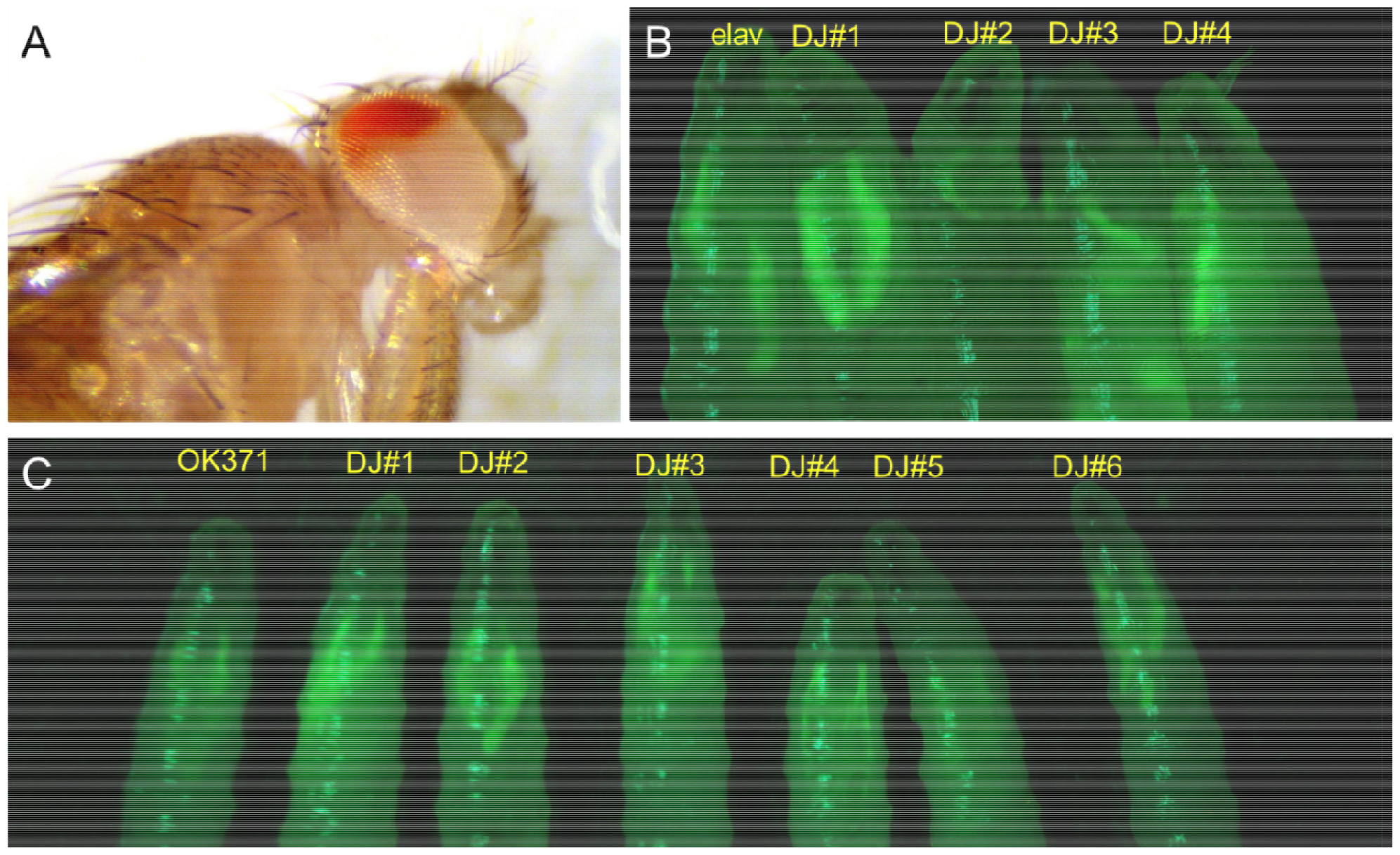
(A) An example of the mosaic-eyed fly, (B) GFP expression in intact larvae with the DIY fluorescence stereoscope setup for elav^C155DJ^ lines, (C) GFP expression in intact larvae with the DIY fluorescence stereoscope setup for OK371^DJ^ lines.

## Notes

### Competing Interest Statement

The authors have declared no competing interest.

